# Novel tau filament fold in corticobasal degeneration, a four-repeat tauopathy

**DOI:** 10.1101/811703

**Authors:** Wenjuan Zhang, Airi Tarutani, Kathy L. Newell, Alexey G. Murzin, Tomoyasu Matsubara, Benjamin Falcon, Ruben Vidal, Holly J. Garringer, Yang Shi, Takeshi Ikeuchi, Shigeo Murayama, Bernardino Ghetti, Masato Hasegawa, Michel Goedert, Sjors H.W. Scheres

## Abstract

Corticobasal degeneration (CBD) is a neurodegenerative tauopathy that is characterised by motor and cognitive disturbances (1–3). A higher frequency of the *H1* haplotype of *MAPT*, the tau gene, is present in cases of CBD than in controls (4, 5) and genome-wide association studies have identified additional risk factors (6). By histology, astrocytic plaques are diagnostic of CBD (7, 8), as are detergent-insoluble tau fragments of 37 kDa by SDS-PAGE (9). Like progressive supranuclear palsy (PSP), globular glial tauopathy (GGT) and argyrophilic grain disease (AGD) (10), CBD is characterised by abundant filamentous tau inclusions that are made of isoforms with four microtubule-binding repeats (4R) (11–15). This distinguishes 4R tauopathies from Pick’s disease, filaments of which are made of three-repeat (3R) tau isoforms, and from Alzheimer’s disease and chronic traumatic encephalopathy (CTE), where both 3R and 4R tau isoforms are found in the filaments (16). Here we report the structures of tau filaments extracted from the brains of three individuals with CBD using electron cryo-microscopy (cryo-EM). They were identical between cases, but distinct from those of Alzheimer’s disease, Pick’s disease and CTE (17–19). The core of CBD filaments comprises residues K274-E380 of tau, spanning the last residue of R1, the whole of R2, R3 and R4, as well as 12 amino acids after R4. It adopts a novel four-layered fold, which encloses a large non-proteinaceous density. The latter is surrounded by the side chains of lysine residues 290 and 294 from R2 and 370 from the sequence after R4. CBD is the first 4R tauopathy with filaments of known structure.

---

We extracted tau filaments from the frontal cortex of three individuals with a neuropathologically confirmed diagnosis of CBD. Abundant neuronal inclusions and astrocytic plaques were stained by antibodies specific for 4R tau (Figure 1a-c) and for hyperphosphorylated tau (Figure 1e), as well as by Gallyas-Braak silver (Figure 1f). Antibodies against 3R tau failed to give specific staining (Figure 1d). By immunoblotting of sarkosyl-insoluble fractions, two major tau bands of 64 and 68 kDa were stained by an antibody specific for 4R tau, as were two minor bands of 37 kDa (Figure 1g). Immunogold negative-stain EM with antibodies specific for the N- and C-termini of tau, as well as for R1, R2, R3 and R4, indicated that epitopes of R2, R3 and R4 form part of the ordered cores of filaments in all three cases of CBD (Extended Data Figure 1). This is consistent with the estimated lengths of trypsin-resistant cores of CBD filaments (20). Narrow and wide filaments were present (Figure 1h), in agreement with previous findings (21). Narrow filaments have a helical twist with a crossover distance of approximately 900 Å and a minimal width of 80 Å, a maximal width of 130 Å. Wide filaments have a crossover distance of approximately 1,300 Å and a maximal width of 260 Å with similar minimal width as narrow ones. We named these filaments Type I (narrow) and Type II (wide) CBD filaments, respectively. The ratios of Type II to Type I filaments ranged from 3:1 to 1:1, depending on cases. Co-pathologies are often found in CBD (22, 23). Small amounts of assembled TDP-43 were present in frontal cortex of CBD cases 1 and 2; CBD case 3 was negative (Extended Data Figure 2). It has been reported that *C9orf72* intermediate repeat expansions are associated with a subset of cases of CBD (24). Such expansions were not present in CBD cases 1-3. CBD case 1 had 2 repeats on each allele, CBD case 2 also had 2 repeats on each allele and CBD case 3 had 2 repeats on one allele and 11 repeats on the other. Staining for Aβ in frontal cortex was stage A for CBD case 1 and stage 0 for CBD cases 2 and 3. Abundant α-synuclein inclusions were not present in CBD cases 1-3, nor were inclusions positive for FUS or positive for dipeptide repeats.

**Figure 1.**
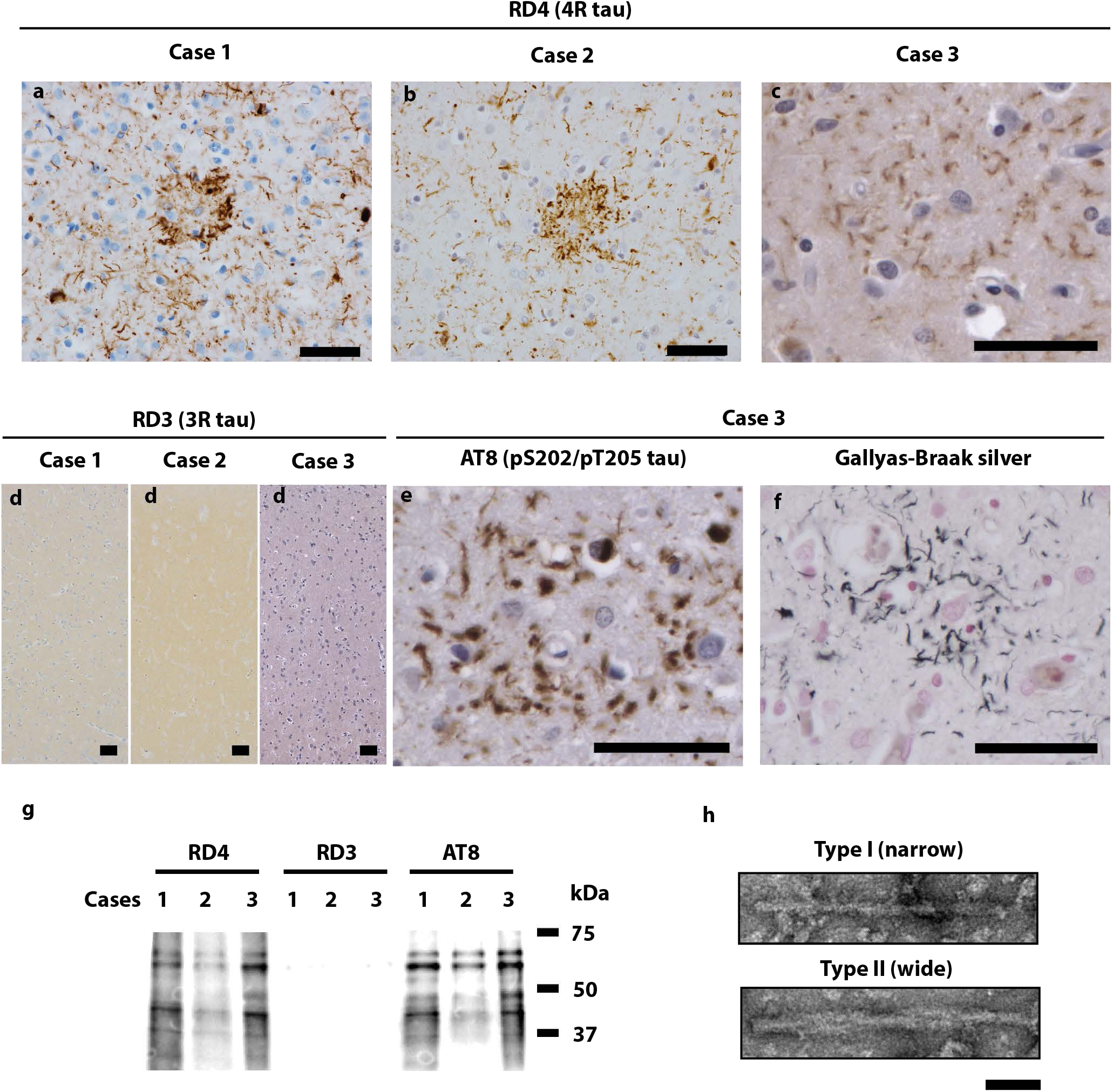
Filamentous tau pathology of CBD. (a-f), Staining of neuronal inclusions, neuropil threads and astrocytic plaques in the frontal cortex of CBD cases 1-3 by antibody RD4 (specific for 4R tau, brown) (a-c), and in the frontal cortex of case 3 by antibody AT8 (pS202, pT205 tau, brown) (e) and Gallyas-Braak silver (black) (f). Staining of frontal cortex from CBD cases 1-3 was negative when antibody RD3 (specific for 3R tau) was used (d). Nuclei were counterstained in blue. Scale bars, 50 μm. (g), Immunoblots using antibodies RD4, RD3 and AT8 of sarkosyl-insoluble tau extracted from the frontal cortex of CBD cases 1-3. (h), Negative-stain electron micrographs of Type I (narrow) and Type II (wide) tau filaments extracted from the frontal cortex of CBD case 1. Scale bar, 50 nm.

We used cryo-EM and helical reconstruction in RELION (25) to determine the structures of both types of tau filaments of CBD (Figure 2a, Extended Data Table 1). In the three cases, Type I filaments are composed of a single protofilament and adopt a novel four-layered fold. Like CTE filaments, each protofilament of CBD contains an additional density that is surrounded by density of the tau protein chain. Unlike CTE (19), the additional density is found in a positively charged environment. Type II filaments consist of pairs of identical protofilaments of Type I, related by C2 symmetry, with less well-resolved maps at the ends of the cores than in their central parts. For case 1, we obtained maps of Type I and Type II tau filaments at overall resolutions of 3.2 Å and 3.0 Å, respectively (Extended Data Figure 3a-d). Local resolution in the central part of Type II filaments extended to 2.8 Å. The maps showed side chain densities and β-strands that were well separated along the helical axis, allowing us to generate stereochemically refined atomic models (Figure 2b, Extended Data Figure 3e-h).

**Figure 2.**
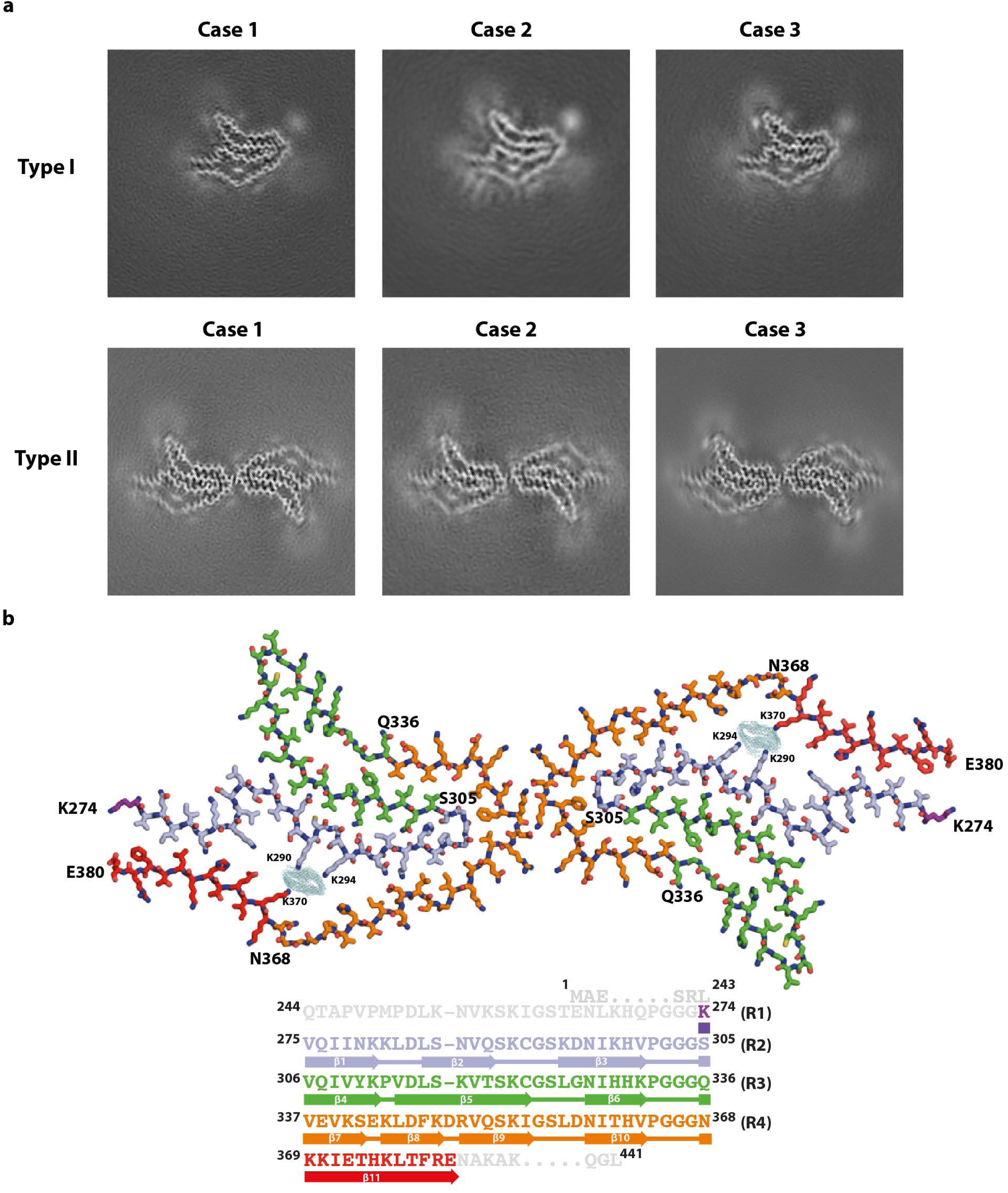
Cryo-EM maps of CBD Type I and Type II tau filaments and atomic model of Type II filaments. (a), Cryo-EM maps of Type I tau filaments (upper panels) and Type II tau filaments (lower panels) from the frontal cortex of cases 1-3. (b), Atomic model of the CBD Type II tau filament (upper panel). The extra density is shown in light blue, with K290, K294 and K370 indicated. Schematic depicting the microtubule-binding repeats (R1-R4) of tau and the sequence after R4 that is present in the core of CBD filaments (all shown in different colours) (lower panel). The positions of β-strands (β1-β11) are indicated.

**Figure 3.**
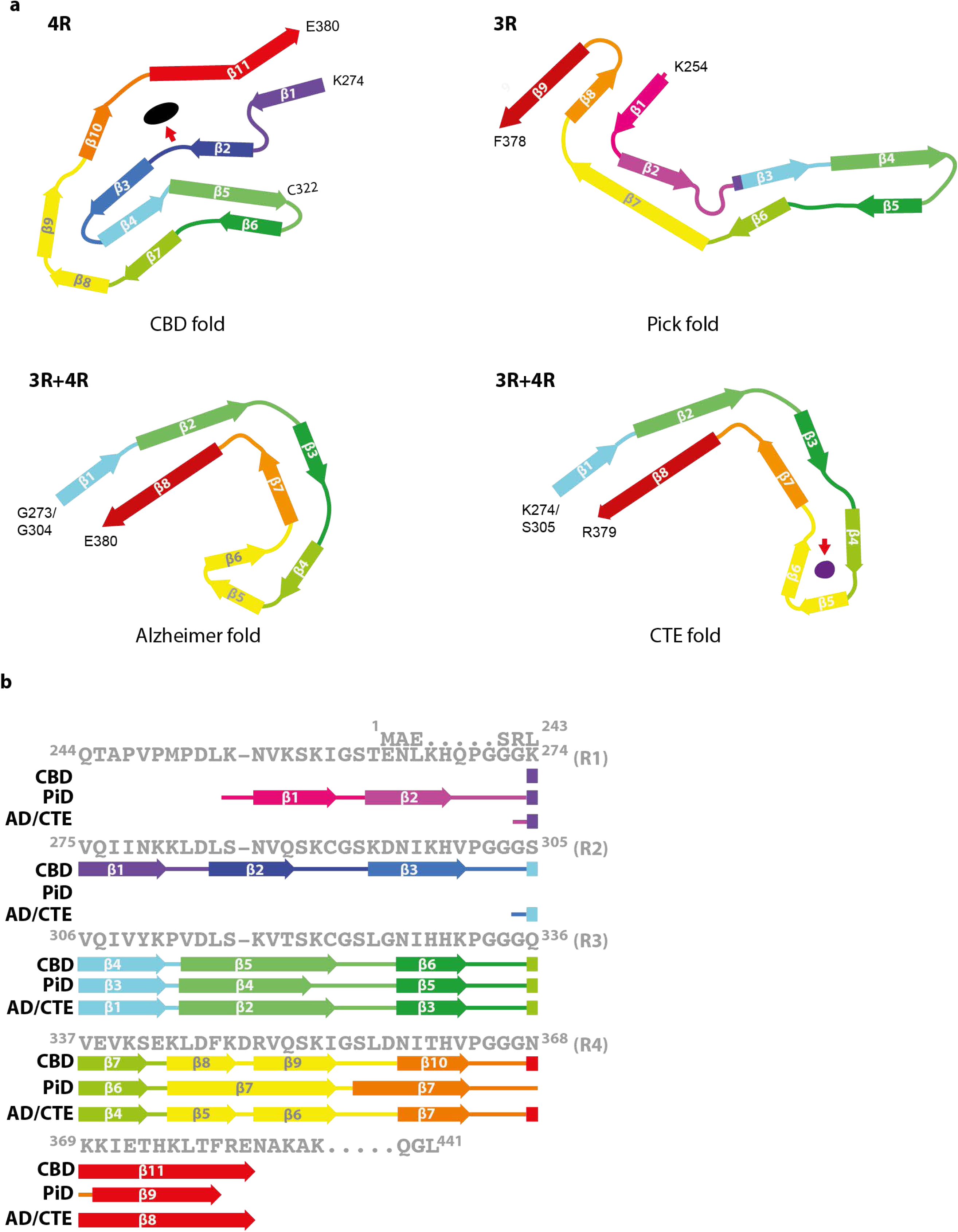
Structures of tau filament cores from human brain. (a), Protofilament from corticobasal degeneration (CBD fold), a 4R tauopathy; protofilament from Pick’s disease (Pick fold), a 3R tauopathy; protofilaments from Alzheimer’s disease (Alzheimer fold) and chronic traumatic encephalopathy (CTE fold), both 3R + 4R tauopathies. Red arrows point to the internal, non-proteinaceous densities in CBD and CTE folds. (b), Schematic depicting the microtubule binding repeats (R1-R4) of tau and the sequence after R4, with the β-strands found in the cores of tau filaments in the different diseases marked by thick arrows.

Type I and Type II tau filaments contain a common protofilament, whose core structure (CBD fold) is composed of residues K274-E380, i.e. the last residue of R1, all of R2-R4, and 12 amino acids after R4. In the core, there are eleven β-strands (β1-β11): three from R2 (β1-β3), three from R3 (β4-β6), four from R4 (β7-β10) and one from the sequence after R4 (β11). They are connected by turns and arcs and form a four-layered structure (Figure 2b, Extended Data Figure 4). The central four layers are formed by β7, β4, β3 and β10. Strands β3 and β4 are connected by a sharp turn, whereas β7 and β10 are connected through β8 and β9, which wrap around the turn. On the other side, β2, β5 and β6 form a three-layered structure. β2 packs against one end of β5 and β6 packs against the other end. The first and the last strands, β1 and β11, pack against each other and close a hydrophilic cavity formed by residues from β2, β3, β10, β11 and the connections between β1 and β2, as well as between β2 and β3 (Figure 2b, Extended Data Figure 4). All interfaces in the CBD fold have mixed compositions of polar and hydrophobic groups (Extended Data Figure 4a).

Each tau repeat contains a PGGG motif (16). In the CBD fold, the PGGG motif of R2 (residues 301 to 304) forms a tight turn between β3 and β4, which is essential for the formation of the four-layered cross-β packing. The PGGG motif of R3 (residues 332 to 335) adopts an extended conformation between β6 and β7, compensating for the shorter lengths of these strands compared to the opposing β4 and β5 connected by P312. The PGGG motif of R4 (residues 364-adopts a similar extended conformation, forming part of the hydrophilic cavity.

In CBD Type II tau filaments, the two protofilaments are related by C2 symmetry (Figure 2b, Extended Data Figure 4). The interface between protofilaments is formed by anti-parallel stacking of ^343^KLDFKDR^349^. Besides van der Waals interactions between the anti-parallel side chains of K347 from each protofilament, the ε-amino group of K347 forms hydrogen bonds with the carboxyl group of D348 and the backbone carbonyl of K347 on the opposite protofilament (Figure 2b, Extended data Figure 5).

The hydrophilic cavity contains an additional density that is different from tau (Figure 2b, Extended Data Figure 4). The positively charged side chains of K290, K294 and K370 point to this extra density, which is not connected to the density for tau, indicating that it is not covalently linked to tau, and therefore cannot represent a post-translational modification. The extra density is as strong as that of tau, suggesting near-stoichiometric occupancy (Figure 2a). In cross-section, the density has approximate dimensions of 9 Å by 4 Å, while it has only few features along the helical axis. Attempts to resolve additional features by performing refinements with a larger helical rise failed. We hypothesise that the extra density is made of non-proteinaceous, polyanionic molecules with a charge of -3 per rung. The buried nature of the negatively charged molecules, their high occupancy and presence in most filaments at end-stage disease indicate that they are continuously incorporated during filament formation. It is therefore possible that these molecules stabilise the CBD fold during initial filament assembly and/or subsequent seeded aggregation. Candidate molecules include cellular metabolites with phosphate and/or carboxylic groups, such as polyphosphate and phosphoglycerate.

CBD is characterised by abundant neuronal and glial inclusions of 4R tau. It has been suggested that tau inclusions may form first in astrocytes of striatum and prefrontal cortex (26). Astrocytic plaques, in which assembled 4R tau is present in the end-feet of astrocytes (6), have been reported to have a perivascular localisation (27), suggesting that in CBD cofactors for tau assembly may enter the brain from the periphery, similar to what we have hypothesised for CTE (19).

Six tau isoforms are expressed in adult human brain, three isoforms with 3R and three isoforms with 4R (28). Based on the tau isoform composition of their filaments, three different types of tauopathies can be distinguished: 3R, 4R and 3R+4R. We previously reported the structures of tau filaments from Alzheimer’s disease, Pick’s disease and CTE (Figure 3) (17–19). Pick’s disease is a 3R tauopathy, whereas both Alzheimer’s disease and CTE are 3R+4R tauopathies. Tau filaments from Alzheimer’s disease and CTE are different, indicating that tau isoform composition is not the sole determinant of conformation. CBD is the first example of a 4R tauopathy of known filament structure. Differences in protofilament structure are observed between diseases, but not between subjects with a given disease, consistent with the existence of distinct conformers of assembled tau in different tauopathies (Figure 3).

The cores of tau filaments from human brain of known structure all contain R3, R4 and 10-12 amino acids after R4 (17–19). Filaments comprising 3R+4R tau in Alzheimer’s disease and CTE do not have R1 or R2 in their cores, and those comprising 3R tau in Pick’s disease have part of R1, whereas filaments comprising 4R tau in CBD contain the whole of R2. This makes the CBD fold the largest known tau fold, with 107 ordered residues. Therefore, filament disassembly may come at a relatively high energetic cost, which may in turn have implications for seeded aggregation and disease progression. Tau assemblies from CBD brains have been shown to seed specific aggregation (29–31). It is likely that filaments from other 4R tauopathies, such as PSP, GGT and AGD, have also at least part of R2 in their cores, but these structures remain to be determined.

It was previously not known why only 4R tau isoforms are present in the filaments of CBD. Our structure reveals that S305, the last residue of R2, which starts β4, is located at a position, where the side chain of K274, the last residue of R1, cannot fit (Figure 2b). Moreover, if R1 were incorporated instead of R2, K294 would be replaced by T263, which would weaken the interaction with the extra density (Figure 2b,d). In support, the sarkosyl-insoluble fraction from CBD cases 1-3, which was used for cryo-EM, seeded aggregation of soluble tau in SH-SY5Y cells expressing full-length 4R, but not 3R, human tau (Extended Data Fig. 6). Similarly, CBD-tau recruited only soluble 4R tau into insoluble aggregates in primary neurons, whereas Alzheimer’s disease-tau recruited both 3R and 4R tau (31). In contrast, tau filaments extracted from the brain of a patient with Pick’s disease seeded aggregation of 3R, but not 4R, human tau (18). Templated misfolding of this type may explain why only 4R tau is incorporated into CBD filaments.

Despite differences between folds, with the structures of tau filaments from four human tauopathies now known, common patterns are beginning to emerge (Figure 3). Packing between β1 and β11 in the CBD fold resembles that between β1 and β8 in the Alzheimer and CTE folds (18, 20). In CBD, ^274^KVQIINK^280^ packs against ^374^HKLTFRE^380^, whereas in Alzheimer’s disease and CTE ^305^KVQIVYK^311^ packs against ^374^HKLTFRE^380^. Hexapeptides VQIINK and VQIVYK are necessary for cofactor-induced assembly of recombinant tau (32, 33), seeded aggregation in cultured cells (34) and assembly of mutant human tau in transgenic mice (35). Residues ^374^HKLTFRE^380^ are missing from the widely used tau constructs K18 and K19, which end at E372 (36). The hairpin-like structure of β4-β7 in the CBD fold resembles that of β3-β6 in the Pick fold, with the exception of C322, which points inwards to form the sharp turn in the CBD fold, and which points outwards in the Pick fold. Interestingly, in all four tau filament folds from human brain, β-strands are formed by approximately the same residues (Figure 3); this is also true of tau filaments assembled *in vitro* using heparin (37). It suggests a model for the diversity of tau folds, where β-strands form fixed building blocks and the loops and turns between strands provide diversity. Tau repeats contain many glycine and proline residues, all of which are located in loops and turns.

This work may shed light on why tau folds differ between diseases, which may in turn reveal mechanisms that lead to ordered assembly. Post-translational modifications may be important. Thus, deamidation of N279 in R2 of tau takes place in Alzheimer’s disease, but not in CBD or PSP (38). This residue, which is located in β1 of CBD, is outside the structured core of AD and CTE filaments. Association of non-proteinaceous cofactors with tau filaments from human brain was unexpected, even though it is well established that such factors can induce assembly of soluble, unmodified tau protein *in vitro* (16). In the same way that hydrophobic molecules inside the β-helix may shape the CTE fold (19), polyanionic molecules inside the positively charged cavity may help to form the CBD fold. We previously speculated that the extra densities near K317 and K321 of tau on the periphery of the Alzheimer fold (17), which are also observed in the CTE fold (19), may be formed by ^7^EFE^9^ of tau. They are believed to be essential for the formation of the straight tau filaments of Alzheimer’s disease. Their similarity to the extra density in the CBD fold raises the possibility that non-proteinaceous molecules may also play a role in providing specificity for tau assembly into Alzheimer and CTE folds.

Determination of the CBD fold opens 4R tauopathies to structural analysis. It supports the hypothesis that distinct conformers of filamentous tau define different tauopathies. We previously showed that tau filaments from Alzheimer’s disease, Pick’s disease and CTE adopt different folds (17–19). Future investigations into what drives the specificity of tau conformers in tauopathies may lead to novel therapeutic opportunities.

## Acknowledgements

We thank the patients’ families for donating brain tissues; F. Epperson, R.M. Richardson and U. Kuederli for brain collection and technical support with neuropathology; G. Cannone, S. Chen, J. Brown, G. Sharov, and A. Yeates for support with electron microscopy; T. Nakane for help with RELION; G. Murshudov and R. Warshamanage for help with REFMAC; P. Emsley for help with COOT; T. Darling and J. Grimmett for help with high-performance computing; R.A. Crowther for helpful discussions. W. Z. was supported by a Foundation that prefers to remain anonymous. M.G. is an Honorary Professor in the Department of Clinical Neurosciences of the University of Cambridge. This work was supported by the U.K. Medical Research Council (MC_U105184291 to M.G. and MC_UP_A025_1013 to S.H.W.S.), the European Union (Joint Programme-Neurodegeneration Research REfrAME, to B.F. and M.G., and the EU/EFPIA/Innovative Medicines Initiative [2] Joint Undertaking IMPRiND, project 116060, to M.G.), the Japan Agency for Science and Technology (Crest, JPMJCR18H3), to M.H., the Japan Agency for Medical Research and Development (JP18ek0109391 and JP18dm020719), to M.H. and (JP19ek0109392), to T.I., the U.S. National Institutes of Health (P30AGO10133 and UO1NS110437), to R.V. and B.G., and the Department of Pathology and Laboratory Medicine, Indiana University School of Medicine, to R.V. and B.G. This study was supported by the MRC-LMB EM facility. We acknowledge the Center for Medical Genomics of Indiana University School of Medicine for next-generation DNA sequencing.

## Author contributions

A.T., K.L.N., T.M., S.M., B.G. and M.H. identified patients, performed neuropathology and extracted tau filaments from CBD cases 1 and 2; R.V., H.J.G. and T.I. carried out genetic analyses; W.Z. extracted tau filaments from CBD case 3 and conducted immunolabelling. W. Z. and B. F. performed cryo-EM; W.Z., Y.S. and S.H.W.S. analysed the cryo-EM data; W.Z. and A.G.M. built the atomic models; A.T. and M.H. carried out seeded aggregation; M.G. and S.H.W.S. supervised the project; all authors contributed to writing the manuscript.

## METHODS

### Clinical history and neuropathology

Frontal cortex from three patients with a neuropathologically confirmed diagnosis of CBD was used. Case 1 was a female from Japan who died aged 74 following a six year history of progressive memory loss and motor impairment. Case 2 was a female from Japan who died aged 79 following a nine year history of progressive memory impairment and spatial disorientation. Case 3 was a male from the U.S. who died aged 52 following a seven year history of personality changes and cognitive dysfunction. Neuropathologically, the brains of all three cases exhibited signs of CBD, with abundant 4R tau- and silver-positive neuronal inclusions, neuropil threads and astrocytic plaques (Figure 1a-e). Tufted astrocytes characteristic of progressive supranuclear palsy (PSP) were not observed. By immunoblotting of the sarkosyl-insoluble fraction, bands of 64 and 68 kDa were detected by antibodies RD4 and AT8, but not by RD3. Bands of 37 kDa were also seen, consistent with CBD (Figure 1g). No known disease-causing mutations were present in *MAPT*. All three cases were homozygous for A at position 152; A152T tau has been reported to be a risk factor for some tauopathies (39, 40). Whole-exome and whole-genome sequencing did not detect mutations known to cause Alzheimer’s disease, Parkinson’s disease, frontotemporal dementia or amyotrophic lateral sclerosis. All three cases were homozygous for the *H1* haplotype of *MAPT*. Two haplotypes, *H1* and *H2*, are present in the Caucasian population, with the *H1* haplotype being over-represented in individuals with CBD and PSP (4, 5). The Japanese population has only *H1* (41, 42). The *APOE* genotypes of CBD cases 1-3 were: case 1 (ε3/ε4), case 2 (ε3/ε3), case 3 (ε3/ε3).

### Whole-exome sequencing

Target enrichment made use of the SureSelectTX human all-exon library (V6, 58 Mb, Agilent) and high-throughput sequencing was carried out using a HiSeq4,000 (2×75-bp paired-end configuration, Illumina). Bioinformatics analyses were performed as described (43).

### Whole-genome sequencing

Sequencing libraries were prepared using 100 ng high-quality genomic DNA from cerebellum using Illumina Nextera DNA Flex Library Prep Kit, and assessed using a Qubit and Agilent Bioanalyzer. High-throughput DNA sequencing was carried out on multiple libraries pooled in equal molarity using a NovaSeq6,000 (150-bp paired-end configuration, Illumina) and aligned to the human reference genome GRCh38 using BWA and Bwakit (v.0.7.15). ExpansionHunter was applied to estimate expansion numbers of short tandem repeats (v.2.5.5) and germline variants were identified with strelka2 (v.2.9.9), with default parameters for whole-genome sequence data. The variants were annotated for their effects with ANNOVAR (44).

### *C9orf72* hexanucleotide repeat expansion

Repeat-primed polymerase chain reaction was used to determine the number of GGGGCC hexanucleotide repeats in the first intron of *C9orf72* (Asuragen AmplideX PCR/CE *C9orf72* kit). Internal standards were analysed along with samples to evaluate assay performance. Repeat numbers of up to 25 repeats were determined with an accuracy of ±1 repeat, and repeat numbers greater than 25 were determined with an accuracy of ±3 repeats.

### Extraction of tau filaments

Sarkosyl-insoluble material was extracted from fresh-frozen frontal cortex of CBD cases 1-3, essentially as described (20). Briefly, tissues were homogenised in 20 volumes (v/v) extraction buffer consisting of 10 mM Tris-HCl, pH 7.5, 0.8 M NaCl, 10% sucrose and 1 mM EGTA. Homogenates were brought to 2% sarkosyl and incubated for 30 min. at 37°C. Following a 10 min. centrifugation at 20,000 g, the supernatants were spun at 100,000 g for 20 min. The pellets were resuspended in 700 μl/g extraction buffer and centrifuged at 9,500 g for 10 min. For CBD cases 1 and 2, the supernatants were diluted 3-fold in 50 mM Tris-HCl, pH 7.5, containing 0.15 M NaCl, 10% sucrose and 0.2% sarkosyl and spun at 166,000 g for 30 min. For CBD case 3, the supernatant was spun at 100,000 g for 60 min and the pellet resuspended in 700 μl/g extraction buffer and centrifuged at 9,800 g. The supernatant was then spun at 100,000 g for 60 min. Sarkosyl-insoluble pellets of CBD cases 1-3 were resuspended in 25 μl/g of 20 mM Tris-HCl, pH 7.4, 100 mM NaCl and used for cryo-EM. For immuno-EM the samples were diluted 5-10-fold. Sarkosyl-insoluble pellets of approximately 2 g frontal cortex were used for cryo-EM.

### Immunolabelling, histology and silver staining

Western blotting and immunogold negative-stain EM were carried out as described (45). For Western blotting, the samples were resolved on 4-20% Tris-glycine gels (Novex) and the primary antibodies were diluted in PBS plus 0.2% Tween-20, 1% bovine serum albumin. Primary antibodies were: RD3 and RD4 (46) (Millipore), used at 1:4,000; AT8 (47) (specific for pS202/pT205 tau) (Thermo Fisher), used at 1:1,000 and anti-phospho-TDP-43 (pS409/pS410) (48, 49) (Cosmo Bio), used at 1:1,000. For immuno-EM, primary antibodies were used at 1:50. They were: BR133 (28) (raised against tau residues 1-16), BR136 (18) (raised against tau residues 244-257), Anti-4R (38, 50) (raised against tau residues 275-291), BR135 (28) (raised against tau residues 323-335), TauC4 (20) (raised against tau residues 354-369) and BR134 (28) (raised against tau residues 428-441). Histology and immunohistochemistry were carried out as described (51). Brain sections were 8 micron-thick and were counterstained with haematoxylin. Primary antibodies were: RD3 (1:1,000); RD4 (1:1,000); AT8 (1:300); anti-pS396 tau (1:1,000, Calbiochem); anti-TauC, specific for tau at residues 422-438 (1:1,000, Cosmo Bio); anti-phospho-TDP-43 (1:1,000); anti-FUS (1:200, Sigma); anti-poly-GA (1:1,000, Cosmo Bio); anti-α-tubulin (1:1,000, Sigma) and anti-influenza hemagglutinin (HA) (1:1,000, Sigma). Sections were silver-impregnated using the method of Gallyas-Braak to visualise inclusions (52, 53).

### Seeded tau aggregation

Seeded aggregation was carried out as described (54), except that sarkosyl-insoluble tau from CBD cases 1-3 was used as seed. Briefly, 10 ng sarkosyl-insoluble tau seed and Multifectam (Promega) were added to SH-SY5Y cells transiently expressing HA-tagged 1N3R or 1N4R human tau. Mock transfections were done in the absence of tau seeds. After three days of culture, sarkosyl-insoluble and sarkosyl-soluble fractions were prepared and used for immunoblotting. Insoluble tau was detected with anti-HA and anti-pS396-tau antibodies. Total tau was detected with anti-TauC. Tau concentrations in the seeds from frontal cortex of CBD cases 1-3 were determined using the Tau ELISA kit Wako (FUJIFILM). The intensity of HA-positive bands in the sarkosyl-insoluble fractions was quantified using Image J software.

### Electron cryo-microscopy

Extracted tau filaments were centrifuged at 3,000 g for 30 s, before being applied to glow-discharged holey carbon grids (Quantifoil Au R1.2/1.3, 300 mesh) and plunge-frozen in liquid ethane using a Thermo Fischer Vitrobot Mark IV. Images were acquired on a Gatan K2-Summit detector in counting mode using a Thermo Fischer Titan Krios microscope at 300 kV. A GIF-quantum energy filter (Gatan) was used with a slit width of 20 eV to remove inelastically scattered electrons. Further details are given in Extended Data Table 1.

### Helical reconstruction

Movie frames were gain-corrected, aligned, dose-weighted and then summed into a single micrograph using MOTIONCOR2 (54). The micrographs were used to estimate the contrast transfer function (CTF) using Gctf (55). All subsequent image-processing steps were performed using helical reconstruction methods in RELION 3.0 (25,56,57). Both types of filaments were selected manually in the micrographs, and the resulting data sets were processed independently. For Type I (narrow) filaments, 135,646 segments were extracted with an inter-box distance of 14.1 Å and a box size of 920 pixels. Initial reference-free 2D classification was performed with images that were downscaled to 230 pixels to speed up calculations. Segments contributing to suboptimal 2D class averages were discarded. Assuming a helical rise of 4.75 Å, a helical twist of −0.9° was estimated from the crossover distance of filaments in the micrographs. Using these parameters, an initial 3D reference was reconstructed from the 2D class averages *de novo*. We then re-extracted the selected segments without downscaling them, and with a smaller box of 330 pixels. Using these segments and the *de novo* initial model low-pass filtered to 15 Å, we carried out 3D auto-refinement. We then used the refined reconstruction, low-pass filtered to 15 Å, as reference for a 3D classification without further image alignment. The segments contributing to the best 3D class were used for subsequent 3D auto-refinement of 24,073 selected segments. Refinement of the helical parameters converged onto a helical twist of −0.845° and a helical rise of 4.786 Å. After Bayesian polishing and CTF refinement, the reconstruction was sharpened with a B-factor of −26.23 Å^2^ (Extended Data Table 1) using the standard post-processing procedure in RELION. The overall resolution of the final map was estimated as 3.2 Å from Fourier shell correlations at 0.143 between the two independently refined half-maps, using phase-randomisation to correct for convolution effects of a generous, soft-edged solvent mask that extended to 30% of the height of the box (58). Local resolution estimates were obtained using the same phase-randomisation procedure, but with a soft spherical mask that was moved over the entire map. For Type II (wide) filaments, 129,812 segments were extracted with an inter-box distance of 14.1 Å and a box size of 460 pixels, which were downscaled to 230 pixels for reference-free 2D classification and particle selection. An initial 3D reference was again reconstructed from the 2D class averages *de novo* by assuming a helical rise of 4.75 Å and a helical twist of −0.65°, based on the estimated cross-over distance of filaments. The resulting reconstruction suggested the presence of two-fold symmetry in the structure. To confirm whether this symmetry also had a translational component, we performed two 3D auto-refinements, one with C2 symmetry and one with C1 symmetry, but imposing a pseudo-2_1_ screw axis. By comparing the results of the refinements, we found that the map obtained following the application of C2 symmetry was of higher resolution. The C2 map also showed separation of β-strands along the helical axis, which was absent in the map refined with the screw symmetry. We applied C2 symmetry in all subsequent refinements. Using 3D classification without image alignment, the segments contributing to the best 3D class were selected and used for 3D auto-refinement with optimisation of helical twist and rise. A final 3D auto-refinement of 20,752 selected segments in boxes of 330 pixels without downscaling converged onto a helical twist of −0.61° and the helical rise at 4.786 Å. Following Bayesian polishing (59) and CTF refinement, the overall resolution of the final map was estimated as 3.0 Å. For Type I and Type II tau filaments in CBD case 2 and CBD case 3, the dataset processing steps were similar to Type I and Type II tau filaments in CBD case 1, with the initial 3D references reconstructed *de novo* independently for each dataset.

### Model building and refinement

The models of the cores of Type I and Type II filaments were built *de novo* in combination of both 3.2 Å and 3.0 Å resolution reconstructions from CBD case 1 using COOT (60). We started model building from the ^301^PGGG^304^ motif, with the aromatic side chains of H299 and Y310 not far away, and worked our way towards the N- and C-terminal regions by manually adding amino acids and by targeted real-space refinement in the high-resolution core part of Type II filaments. Tracing of the chain was confirmed by the fitting of the ^332^PGGG^335^ motif, which neighbours the side chains of H329 and H330. Since the density of side chains of N368-E380 was weak in the 3.0 Å reconstruction of Type II filaments, we assigned the side chains in this region following the 3.2 Å reconstruction of Type I filaments. The structure of one protofilament from Type II filaments was rigid-body fitted into the reconstruction of Type I filaments to build the model of Type I filaments. Because of the lack of interaction between two protofilaments, the conformations of the side chains of K343 and K347 in Type I filaments were assigned differently from their counterparts in Type II filaments, according to the reconstruction. Each model was then translated to give a stack of three consecutive monomers to preserve nearest-neighbour interactions for the middle chain in subsequent refinements. Because most residues adopted a β-strand conformation, hydrogen-bond restraints were imposed to preserve a parallel, in-register hydrogen bonding pattern in earlier stages of Fourier-space refinements. Local symmetry restraints were imposed to keep all β-strand rungs identical. Side chain clashes were detected using MOLPROBITY (61) and corrected by iterative cycles of real-space refinement in COOT and Fourier-space refinement in REFMAC (62) and PHENIX (63). For each refined structure, separate model refinements were performed against a single half-map, and the resulting model was compared to the other half-map to confirm the absence of overfitting (Extended Data Figure 3a,b). The final models were stable in refinements without additional restraints. Statistics for the final models are shown in Extended Data Table 1.

### Ethical review board and informed consent

The studies carried out at Tokyo Metropolitan Institute of Medical Science and at Indiana University were approved through the ethical review process at each Institution. Informed consent was obtained from the patients’ next of kin.

### Data availability

Cryo-EM maps for CBD case 1 will be deposited in the Electron Microscopy Data Bank upon acceptance of manuscript for publication. Atomic models for CBD Type I and Type II tau filaments will be deposited in the Protein Data Bank upon acceptance of manuscript for publication. Whole-exome and whole-genome sequencing data and repeat-primed polymerase chain reaction C9orf72 hexanucleotide repeat expansion data will be deposited in the European Genome-phenome Archive upon acceptance of manuscript for publication.

## EXTENDED DATA FIGURE LEGENDS

**Extended Data Figure 1.**
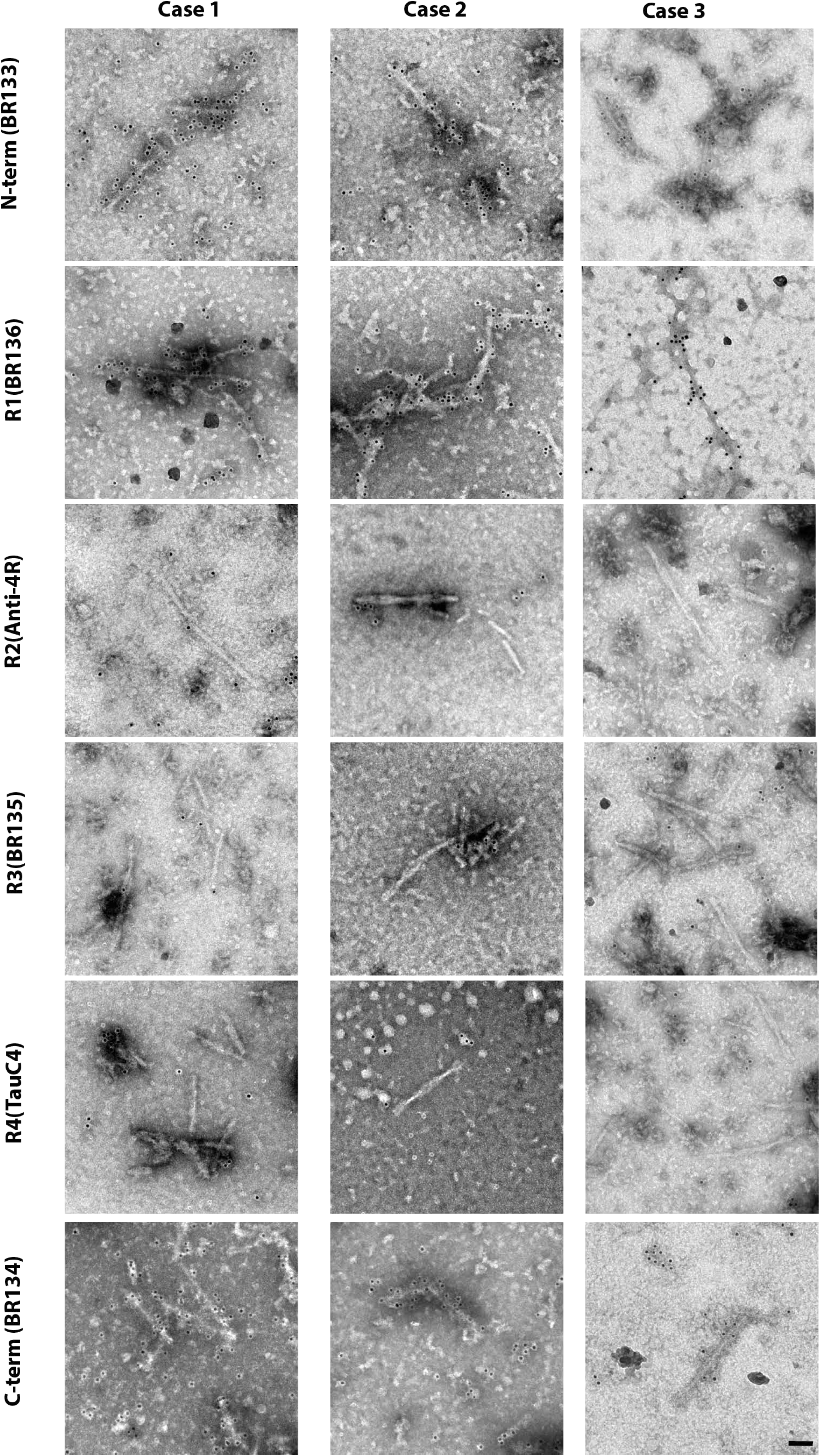
Immunolabelling of tau filaments extracted from the frontal cortex of CBD cases 1-3. Representative immunogold negative-stain electron microscopy images of Type I and Type II tau filaments labelled with antibodies BR133, BR136 and BR134. Antibodies Anti-4R, BR135 and TauC4 did not label filaments, which indicates that their epitopes lie within the ordered filament cores. Scale bar, 50 nm.

**Extended Data Figure 2.**
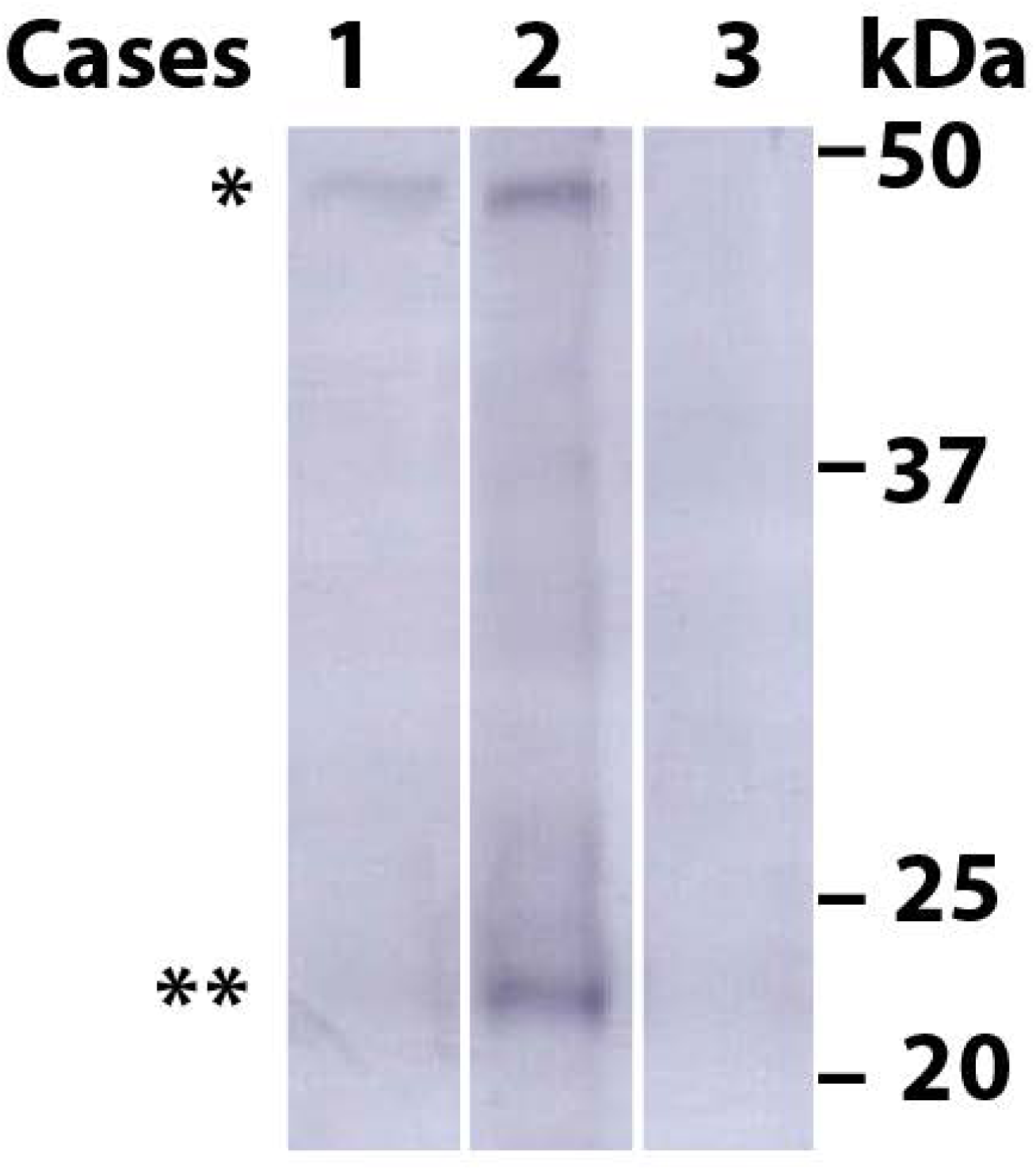
Assembled TDP-43 in frontal cortex of CBD cases 1-3. **a**, Immunoblots using anti-phospho-TDP-43 antibody. Sarkosyl-insoluble material was prepared as described and all the samples were applied on the same gel. The 43 kDa band (*) corresponds to full-length TDP-43 and the 18-26 kDa bands (**) to C-terminal fragments.

**Extended Data Figure 3.**
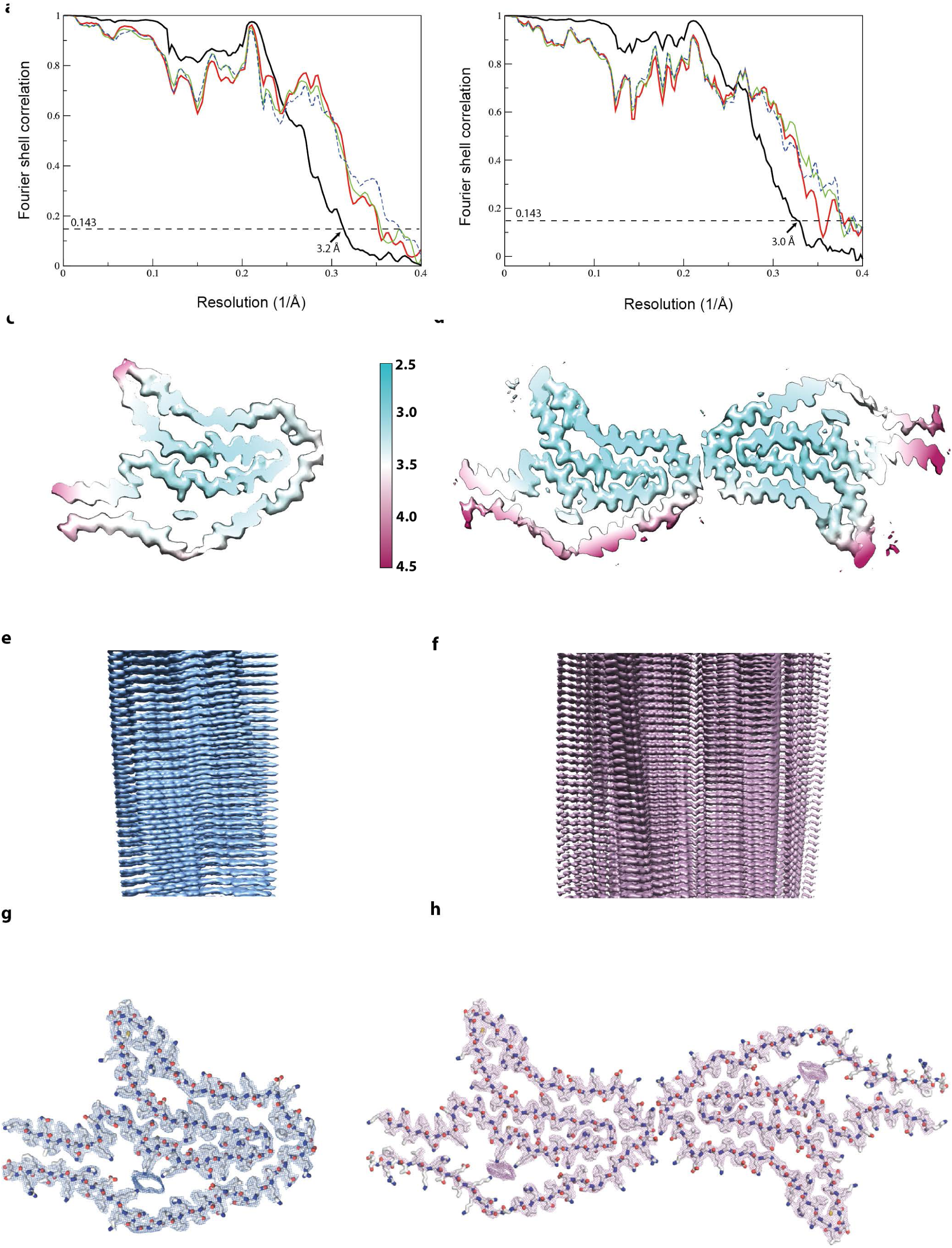
Cryo-EM map and model comparisons. **a, b**, Fourier shell correlation (FSC) curves between two independently refined half-maps (black, solid), of the final model versus the full map (red, solid), of a model refined in the first half-map versus the first half-map (green, solid), and of the same model versus the second half-map (blue, dashed) for CBD Type I (a) and Type II (b) filaments. **c**, **d**, Local resolution estimates for the CBD Type I (c) and Type II (d) filament reconstructions. **e**, **f**, Side views of the 3D reconstructions of CBD Type I (c) and Type II (d) filaments. **g**, **h**, Sharpened, high-resolution cryo-EM maps of CBD Type I (g) and Type II (h) tau filaments with their corresponding atomic models overlaid.

**Extended Data Figure 4.**
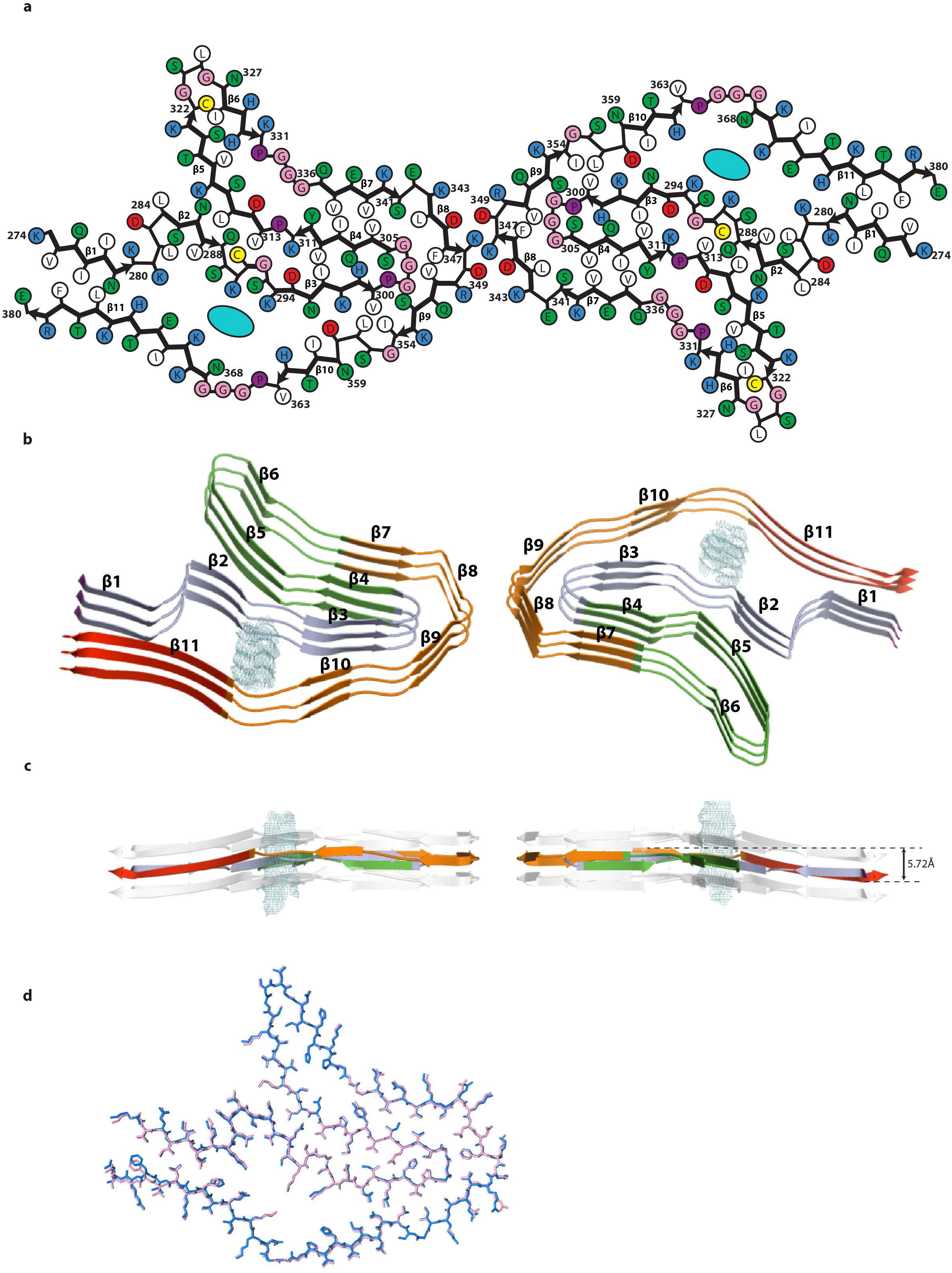
CBD tau filament fold. **a**, Schematic of the CBD fold. **b**, Rendered view of the secondary structure elements in the CBD fold, depicted as three successive rungs. **c**, As in b, but in a view perpendicular to the helical axis, revealing the changes in height within a single molecule. **d**, Comparison of the protofilament structures of CBD Type I (blue) and Type II (pink).

**Extended Data Figure 5.**
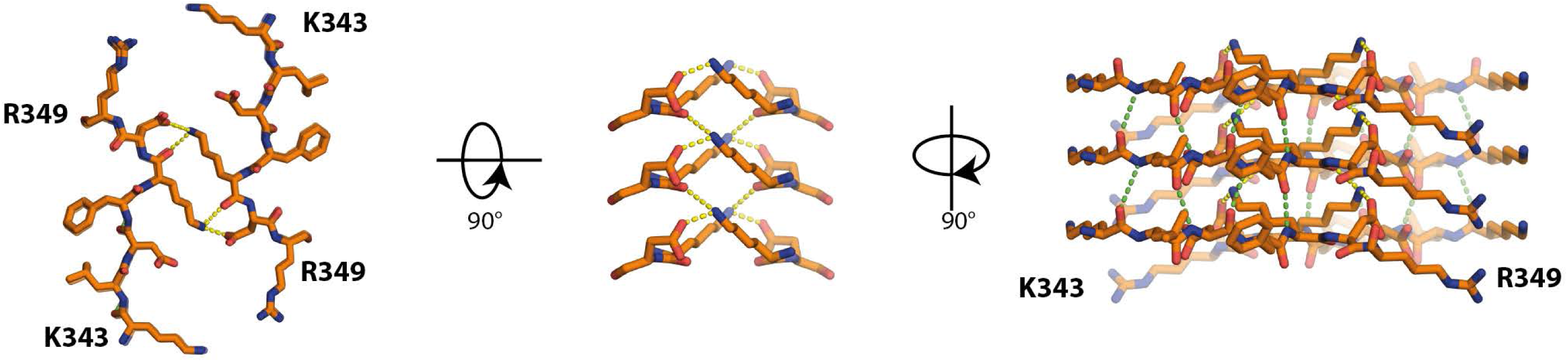
Protofilament interface in CBD Type II tau filaments. Packing between residues ^343^KLDFKDR^349^ of the two protofilaments. Inter-protofilament hydrogen bonds are shown in yellow. Intra-protofilament hydrogen bonds are shown in green.

**Extended Data Figure 6.**
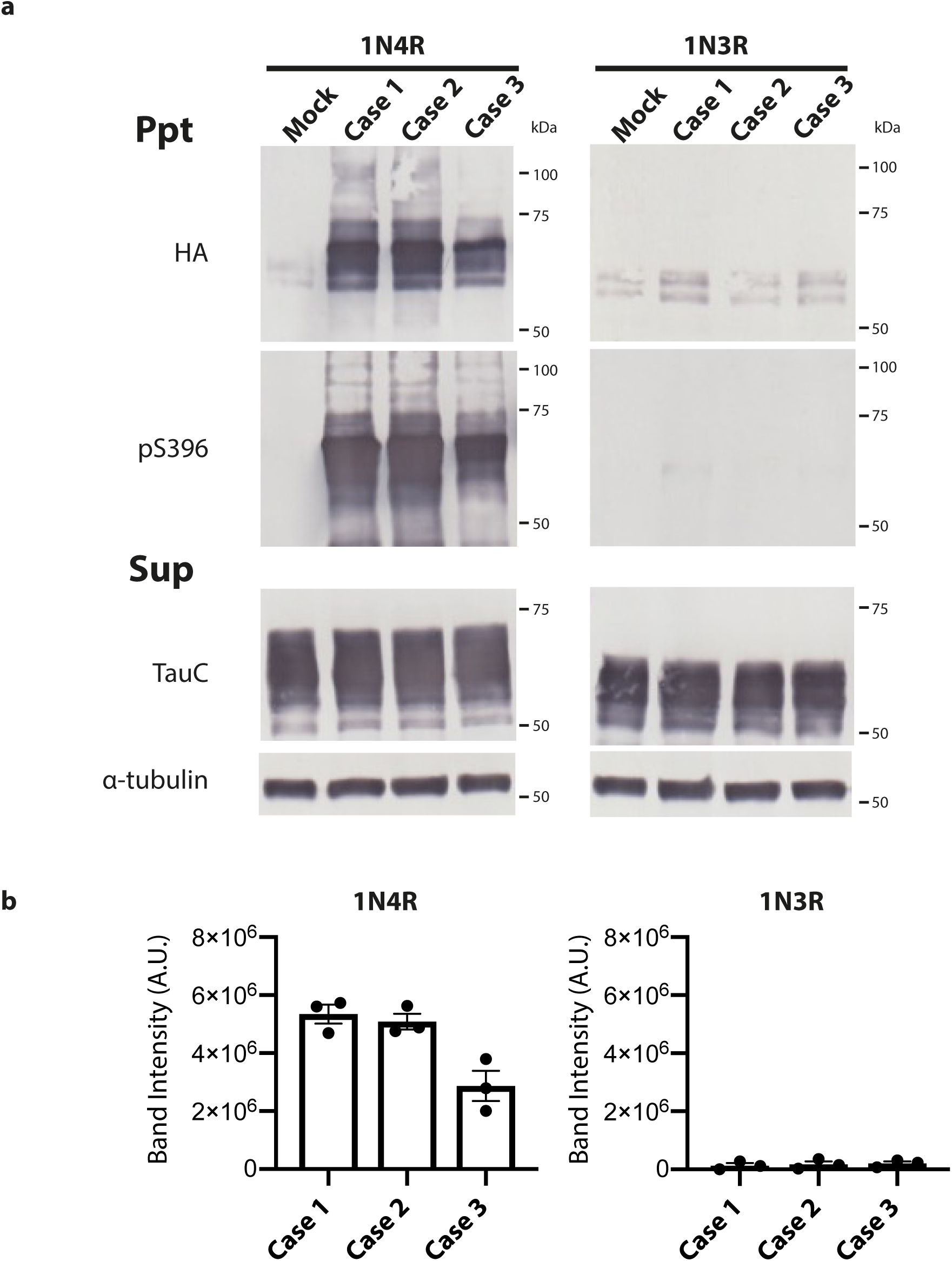
Seeded tau aggregation induced by CBD filaments in SH-SY5Y cells. **a,** Immunoblotting of sarkosyl-insoluble (Ppt) and sarkosyl-soluble (Sup.) fractions extracted from mock-transfected SH-SY5Y cells and from cells transfected with tau seeds from frontal cortex of CBD cases 1-3. SH-SY5Y cells transiently expressed either hemagglutinin (HA)-tagged 1N4R or HA-tagged 1N3R human tau. Insoluble tau was detected with anti-HA and anti-pS396 tau antibodies. Total tau was detected with anti-TauC. Blotting with an anti-α-tubulin antibody served as loading control. **b**, Quantitation of anti-HA-positive bands. The results are expressed as the means ± S.E.M. (n=3).

**Extended Data Table 1.**
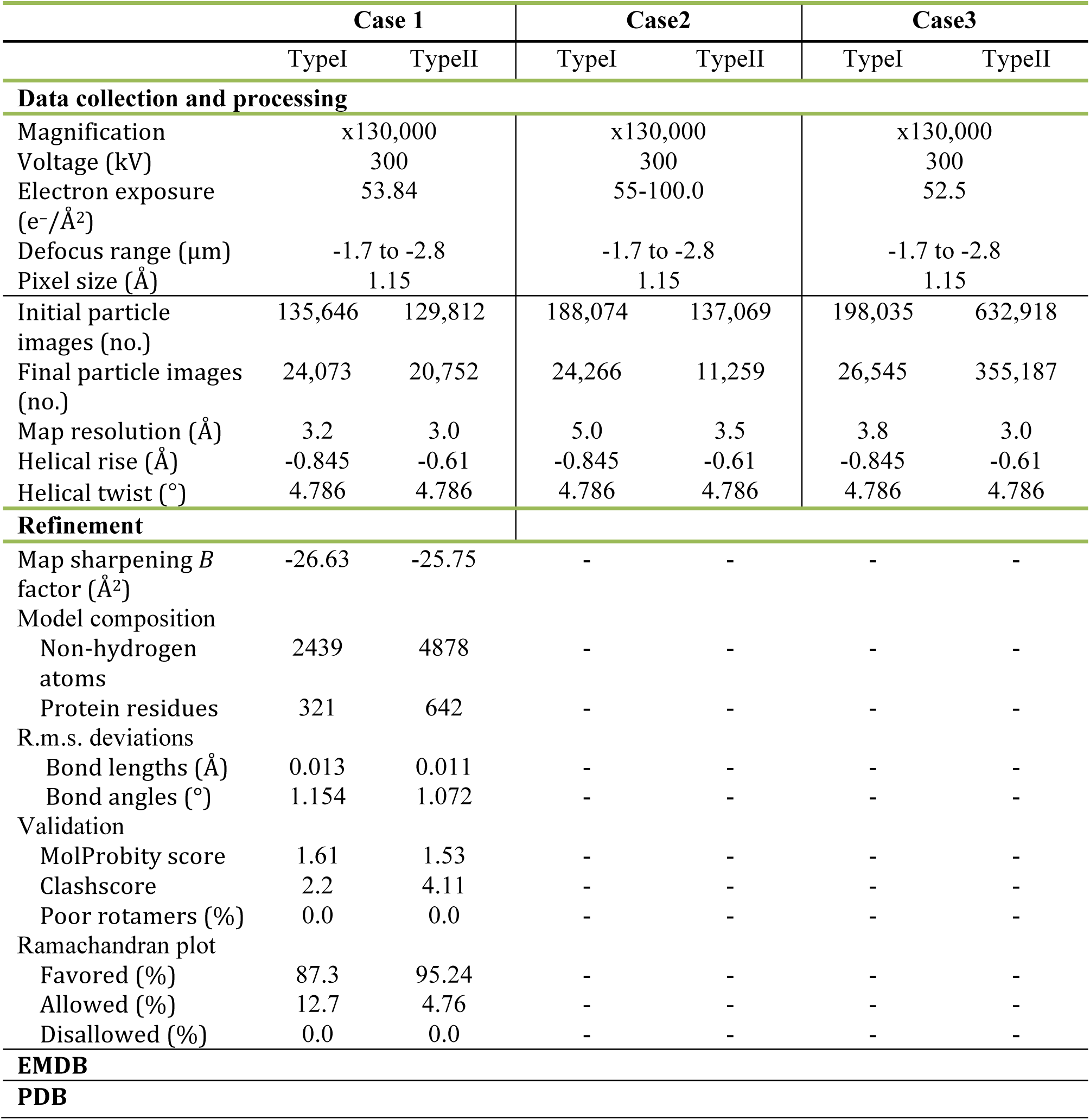
Cryo-EM data collection, refinement and validation statistics.

